# Reduction of fungal dysbiosis is involved in the attenuation of Dextran Sodium Sulfate-Induced Mouse Colitis Mediated by GILZ protein and yeast extract compound

**DOI:** 10.1101/2024.06.18.599634

**Authors:** Marco Gentili, Emilia Nunzi, Samuele Sabbatini, Eleonora Lusenti, Luigi Cari, Antonella Mencacci, Nathalie Ballet, Graziella Migliorati, Carlo Riccardi, Simona Ronchetti, Claudia Monari

## Abstract

Inflammatory bowel diseases (IBD), such as Crohn’s disease and ulcerative colitis, have a complex and multifactorial pathogenesis that remains not fully elucidated. Recent research suggests that intestinal fungal dysbiosis may contribute to the development and persistence of IBD. In this study, we explored, for the first time, the effects of the glucocorticoid-induced leucine zipper (GILZ) protein, known to have protective effects on the gut mucosa in preclinical IBD models, in combination with a yeast extract, which supports the growth of beneficial microorganisms, in a mouse model of ulcerative colitis. The combined treatment produced significant protection against severe disease outcomes in the mice, including the restoration of intestinal barrier integrity and the reduction of pro-inflammatory cytokines. Specifically, GILZ primarily acted on the gut permeability, while the yeast extract mainly reduced pro-inflammatory cytokines. Notably, both treatments were effective in restoring the intestinal burden of clinically important *Candida* and former *Candida* species. Analysis of the intestinal fungal communities revealed that both treatments were able to reduce colitis-associated fungal dysbiosis, promoting a fungal composition similar to that of healthy mice. This effect was mainly the result of a decreased abundance of the *Meyerozima* genus, which was dominant in the colitic mice. Thus, combined treatment regimens with the GILZ protein and yeast extract could represent a new strategy for the treatment of inflammatory bowel diseases, by targeting multiple mechanisms at the basis of IBD, including the fungal dysbiosis.

**IMPORTANCE:** Inflammatory bowel diseases (IBD), including Crohn’s disease and ulcerative colitis, are characterized by chronic inflammation and have a complex, multifactorial pathogenesis that is not yet fully understood. Currently, no established therapeutic strategy can consistently manage IBD effectively. Recent research indicates that intestinal fungal dysbiosis could potentially contribute to the development and persistence of chronic IBD, highlighting the importance of investigating alternative therapeutic strategies able to attenuate fungal dysbiosis in the context of intestinal inflammation. In this study, we demonstrate that a combination of a recombinant protein (GILZp) and a compound with prebiotic properties could represent a new therapeutic strategy for the treatment of IBD, as it not only decreases inflammation and restores the integrity of the epithelial barrier, but reduces fungal dysbiosis associated with DSS-induced colitis.

## INTRODUCTION

Inflammatory bowel diseases (IBD) encompass two important chronic pathologies, namely Crohn’s disease (CD) and ulcerative colitis (UC). Both diseases are characterized by abdominal pain, diarrhea, weight loss, bloody stools, and an inflammatory state owing to granulocytic infiltration in the lamina propria, and this, in turn, causes the persistent release of pro-inflammatory cytokines (1, 2). IBD is also characterized by cycles of alternating relapse and remission phases, so that they impact and reduce the quality of life of patients. Although the pathogenesis of CD and UC has not been completely elucidated, it is known that genomic predisposition, environmental stimuli, dysbiosis of the intestinal microbiota, and aberrant immune response play pivotal roles in their development. So far, no pharmacological treatment is available for the definitive cure of IBD, and the available options are not suitable for all patients. The most common strategy is to use anti-inflammatory drugs or biological agents against inflammatory targets with potentially significant adverse effects (3). The aim of these treatments is to achieve and prolong clinical remissions.

The continuous research on new pharmacological tools has led to the application of recombinant proteins in experimental mouse models of IBD (4, 5). In particular, glucocorticoid-induced leucine zipper (GILZ)-based recombinant proteins present a potentially efficacious treatment for IBD because of their glucocorticoid-like anti-inflammatory properties (6-8). Additionally, we have recently demonstrated that an exogenously administered GILZ protein can control the permeability of the gut in a mouse model of colitis (9), and that GILZ may act as a secretory protein, being expressed by goblet cells in human specimens (10). Therefore, the potential therapeutic effect of GILZ-based proteins in IBD involves both regulation of the integrity and permeability of the gut, and the maintenance of the homeostasis of immune cells.

Over the last few years, numerous studies have documented that patients with IBD, exhibit significant changes in the composition and functionality of the gut microbial communities (11-13) that result in an imbalance between protective and harmful bacteria. The exact mechanisms by which this intestinal dysbiosis contributes to IBD have not been yet fully understood, but it is believed that alterations in the microbial communities can affect intestinal barrier function, mucosal immune responses, and the production of both metabolites and inflammatory molecules. Until a few years ago, the majority of studies focused on the involvement of the bacterial component of the intestinal microbial communities. Only recently, advancements in sequencing technologies and bioinformatics tools have led to significant progress in the understanding of the composition and function of the intestinal fungal community as well as the association between fungal dysbiosis and intestinal inflammation (14-20)). Indeed, it has been reported that changes in fungal diversity and abundance can contribute to the development or exacerbation of intestinal inflammation in IBD (17, 18, 21-25). Although a specific fungal pattern that can define the intestinal mycobiota in patients with IBD has not yet been identified, several studies have reported an increase in the populations of intestinal *Candida spp*. (mainly *C. albicans*, but also *C. tropicalis*, *C. dubliniensis*, and *C. parapsilosis*), as well as former *Candida spp* including *Debaryomyces hansenii* (ex *C. famata)*, *Nakaseomyces glabrata (*ex *C. glabrata)*, *Kluyveromyces marxianus (*ex *C. kefyr*), *Pichia kudriavzevii (*ex *C. krusei*), *Meyerozyma guillermodii* (ex *C. guilliermondii*) and *Clavispora lusitaniae* (ex. *C. lusitaniae*), both in human patients and in preclinical models of IBD (21, 26-34). These recent findings suggest that future therapeutic interventions targeting fungal dysbiosis could be a promising line of research for IBD management.

Prebiotics, are emerging as promising new treatment strategy for IBD (35, 36). Indeed, there is substantial evidence to suggest their potential benefits through multiple mechanisms of action, including promotion of the growth of beneficial bacteria, such as Bifidobacteria and Lactobacilli, increased production of short-chain fatty acids (SCFA), maintenance of gut barrier integrity, regulation of immune responses, and symptom relief (37-41). While most of the research on prebiotics has historically focused on their impact on bacterial communities in the gut, to the best of our knowledge, no study has specifically investigated the impact of prebiotics on the IBD-associated fungal community.

In light of the potential advantages of prebiotics in patients with IBD, it is important to provide data about their role in the treatment of IBD, especially in combination with molecules that exhibit pharmacological activity.

Given the above considerations, the objective of our study was to examine the potential therapeutic impact of combining a recombinant GILZ protein (GILZp) with a yeast extract compound (Py), as a potential prebiotic, in a murine model of colitis. Our results demonstrated that this treatment combination effectively attenuated dextran sulfate sodium (DSS)-induced colitis by reducing inflammation, restoring epithelial barrier integrity, and ameliorating fungal dysbiosis by reestablishing the abundance of the genus *Meyerozyma*.

## MATERIALS AND METHODS

An extended version of material and methods can be found in the supplemental material.

### Animals

Male C57BL/6 mice (6–8 weeks old) were purchased from ENVIGO (Udine, Italy). The mice were housed under specific pathogen-free conditions and a 12/12-h light/dark cycle and received water and food *ad libitum*. The animal care and experimental procedures have been approved by a research ethics committee at the institution at which the studies were conducted, using protocols approved by the Italian Ministry of Health (approval no. 754/2020-PR). The animal studies are reported according to the ARRIVE guidelines (42).

### Compounds

The candidate prebiotic used in this study (NuCel® 582 MG) is a primary yeast (Py) extract obtained by the autolysis of a selected strain of yeast that was grown on a molasses-based media. It is capable of promoting the growth of lactobacilli and is also suitable for ripening flora. This product was provided by Procelys (Lesaffre).

### DSS-induced colitis

Colitis was induced in mice by adding 3% DSS (TdB Consultancy AB, Uppsala Sweden) to autoclaved drinking water *ad libitum* for 5 days. Treatments are detailed in the extended supplemental material. DAI was evaluated daily in each mouse, and values are expressed as previously published (9).

### RNA extraction

mRNA was extracted from the colons using theRNeasy® Plus Mini Kit (Qiagen), following the manufacturer’s instructions. Briefly, after the addition of RLT buffer and β-mercaptoethanol, the samples were homogenized and subsequently transferred into the "gDNAEliminator spin column". After the addition of 70% ethanol, RNA was recovered with a RNAeasy Mini spin column. The extracted mRNA was purified by contaminating DNA with the Wipeout Buffer. Conversion of total mRNA to cDNA was performed using a High-Capacity cDNA Reverse Transcription Kit (Qiagen, Hilden, Germany).

### Real-time quantitative PCR

RNA expression was detected using probes (Thermo fisher) for IL-1β, TNFα, IL-6, IL-12p40, MUC2, and Claudin-2, (Taqman) whose expression was normalized against the 18S housekeeping gene. For data analysis, the relative expression levels were calculated using the 2^-ΔCt^ method.

The DNA extraction procedures for the quantitative analysis of *Candida* and former *Candida* species and the identification of specific forward and reverse primers have been described in the extended supplemental material files.

### Mycobiome sequencing and analysis

Fresh stool samples were collected on day 10 of the experiment and immediately stored in dry ice. The samples were then processed for DNA extraction using the E.Z.N.A.® Stool DNA Kit (Omega Bio-tek) according to the protocol provided with the kit. DNA purity and quantity were determined with a spectrophotometer, and the DNA samples were prepared for sequencing. The sequencing was outsourced to BMR Genomics (Padova, Italy), as detailed in the extended supplemental material. The analysis was performed as described in the extended supplemental material.

### Statistical analyses

All data are expressed as mean ± standard error. Differences between mean values were tested using the GraphPad Prism 6.0 software. Statistical analyses were performed by one-way analysis of variance (ANOVA) with Tukey’s *post hoc* test. Statistical significance was indicated as *P < 0.05, **P < 0.01, ***P < 0.001, ****P < 0.0001.

## RESULTS

### Amelioration of DSS-induced colitis by combined treatment with GILZp and Py

We recently demonstrated that treatment with a recombinant GILZ protein (GILZp) can alleviate symptoms of DSS-induced colitis by restoring gut permeability (9). This is crucial because a leaky gut epithelium allows microbial communities to translocate across intestinal cells, triggering inflammation as the main cause of IBD. Despite the focus on restoring gut balance (eubiosis) in IBD with probiotics and prebiotics, limited or no beneficial effects have been achieved with prebiotics as adjuvant treatment (43-45). We, therefore, investigated Py, a potential new yeast-extracted prebiotic, combined with GILZp in a mouse model of colitis. Py was administered in escalating doses daily via gavage for 8 days, starting from day 0, with administration ceased for the final 2 days. Despite the susceptibility of these mice to DSS-induced mortality (46), those treated with Py showed improved survival rates compared to DSS control (Ctrl) (Fig. S1). Survival was dose-dependent, with the highest dose (1000 mg/kg) ensuring survival until sacrifice, thus utilized in subsequent experiments.

To investigate the effect of combined treatment with GILZp and Py, mice were randomly allocated into five groups, including one group that comprised non-colitic mice and four groups that comprised mice with DSS-induced colitis. Among the colitic groups, one group was left untreated (Ctrl), one group was treated with only Py, one group was treated with only GILZp, and one group received the combined GILZp-Py treatment. The treatments were administered daily until day 8 (Fig. 1A). During the 10-day follow-up period, we observed that mortality was reduced with both the singular and combined treatments, whereas mortality was recorded from day 8 for the DSS Ctrl group (Fig. 1B). Body weight tended to decrease in all the colitic groups, with no statistically significant difference between the treatment and Ctrl groups (Fig. 1C). In contrast, the disease-associated index (DAI) score was significantly different across groups; in particular, the only-Py and combined GILZp-Py treatment groups had a lower DAI score than the Ctrl group (Fig. 1D). However, on the day of sacrifice (day 10), no differences were observed in either body weight or DAI score. Notably, all treatments resulted in a significant reduction in the colon weight/length ratio in comparison with the Ctrl group, suggesting mild inflammation in the treated groups (Fig. 1E).

**FIG. 1.**
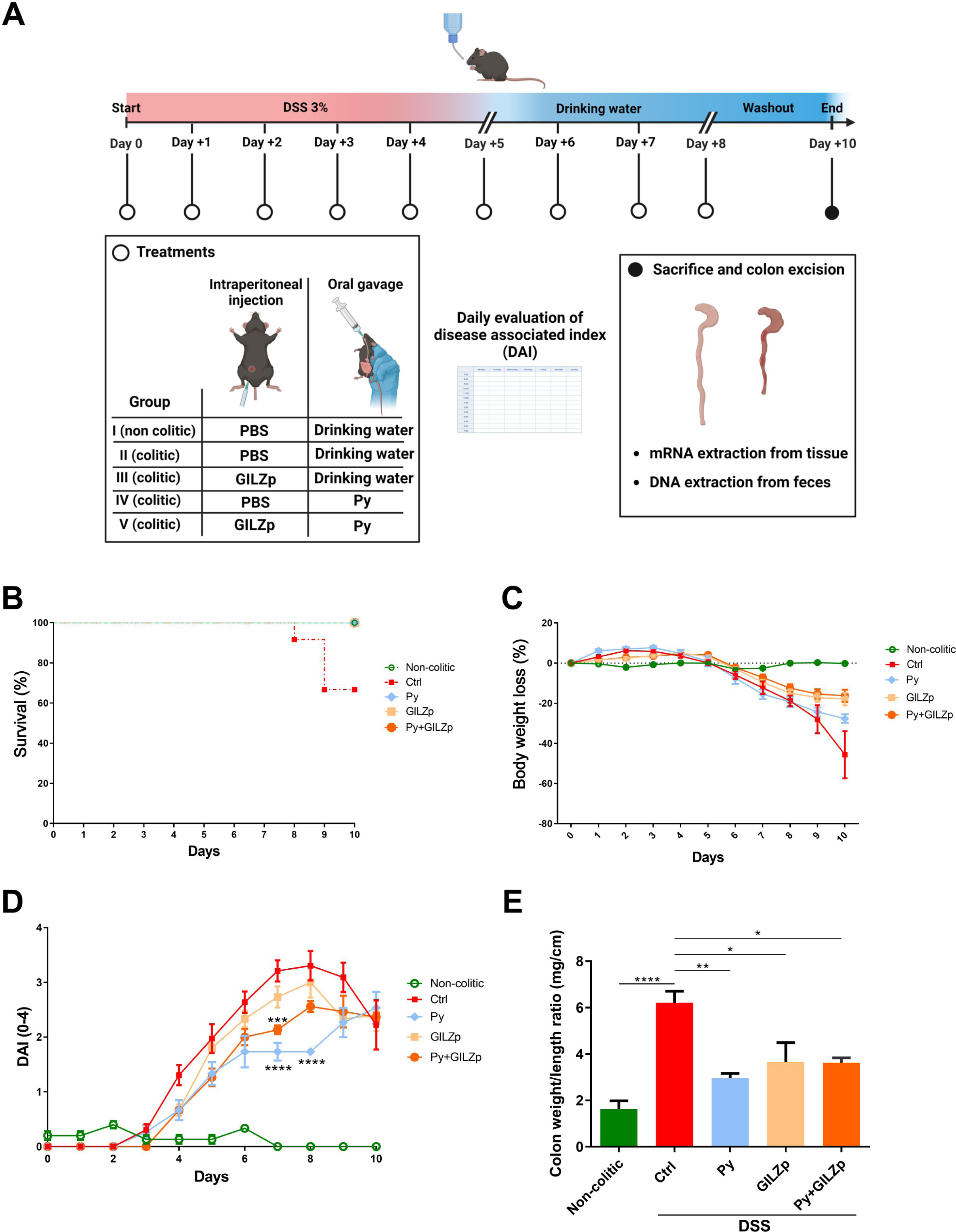
Ameliorative effect of individual and combined GILZp and Py treatments on DSS-induced colitis symptoms in mice. **A** Schematic diagram of the experimental design and procedures. Colitis was induced with the administration of 3% DSS for 5 days. From day 0 to day 8, the four colitic groups received GILZp (0.2 mg/kg), Py (1000 mg/kg), a Py+GILZp combination, or PBS. The schematic was created with Biorender (Biorender.com). **B** Kaplan-Meier curve depicting mortality in mice with DSS-induced colitis. **C** and **D** Body weight loss and clinical score (DAI) were registered daily over the whole experimental period. **E** Mean weight/length ratio values of colons. Values are expressed as the means ± SEM value from two independent experiments (*n* = 5–7 for each group). *P < 0.05, **P < 0.01, ***P < 0.001, ****P < 0.0001

### GILZp and Py restore intestinal barrier integrity and reduce colon inflammation

Since one of the main and more striking observations was the reduction or even absence of mortality when Py and GILZp were used for treatment, we tried to understand the molecular mechanisms underlying this effect. To this end, we analysed the main pro-inflammatory cytokines associated with DSS-induced colitis, such TNFα, IL-1β, IL-12p40, and IL-6. The results showed that while Py was able to suppress the expression of all the inflammatory cytokines, GILZp was only able to suppress IL-1β and TNFα expression and did not affect the other cytokines (Fig. 2A). Further, with the combined treatment, all the pro-inflammatory cytokines, except for IL-12p40, were downregulated: this effect is probably attributable to Py (Fig. 2A).

**FIG. 2.**
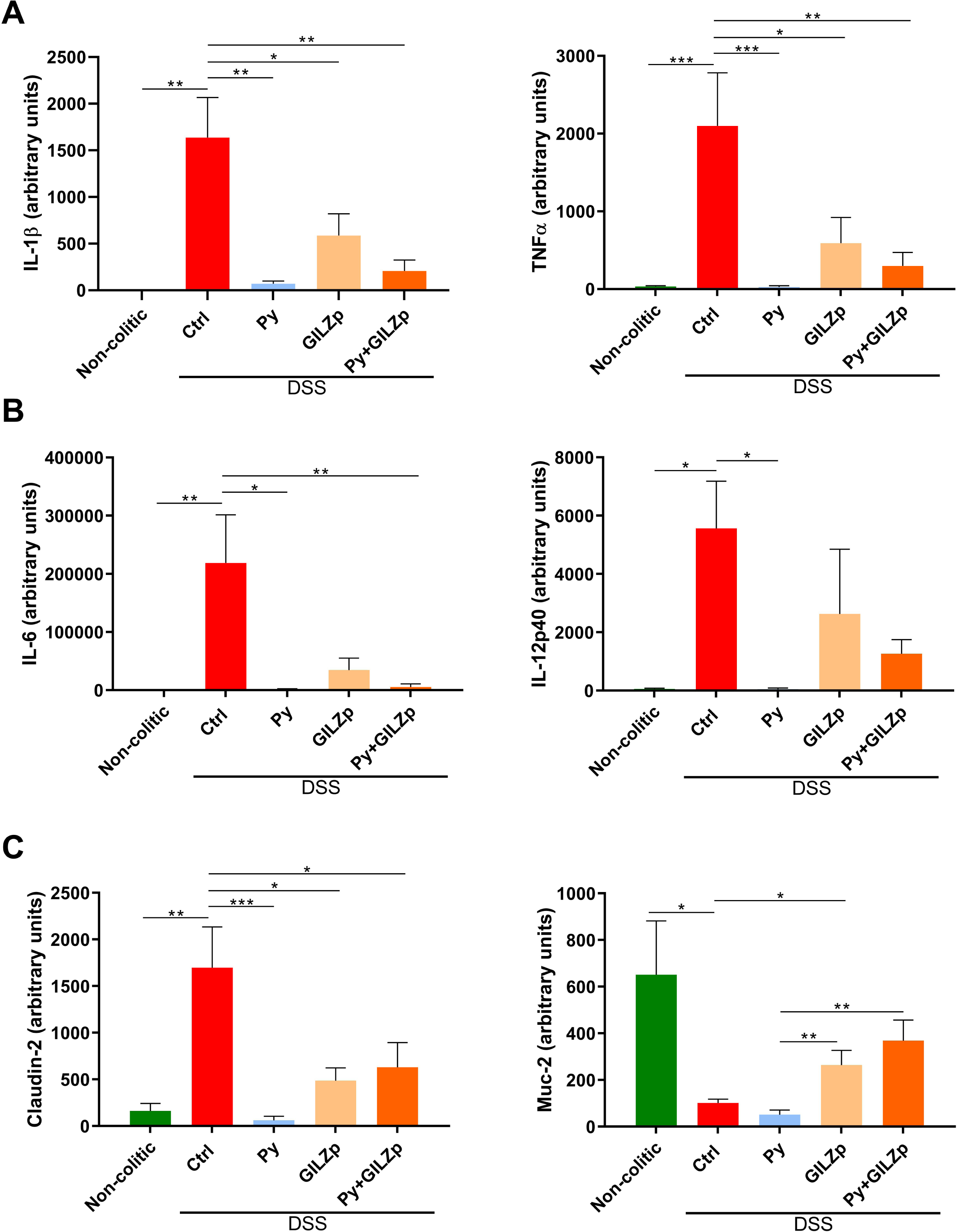
Suppression of the expression of inflammatory cytokines and restoration of intestinal barrier integrity with individual and combined GILZp and Py treatments. Quantitative RT–PCR analysis of IL-1β and TNF-α (**A**) expression, (**B**) IL-6 and IL-12p40 expression and (**C**) Claudin-2 and MUC2 expression in the colon of non-colitic and colitic mice. The colitic mice were treated as indicated. Values are expressed as means ± SEM. (*n* = 5-7) for each group. * *P* < 0.05, ** *P* <0.01, *** *P* <0.001.

In the pathogenesis of colitis, increase in the permeability of the intestinal mucosal barrier and loss of mucosal function are detrimental factors that need to be corrected for mucosal healing. We, therefore, analysed the expression of Claudin-2, one of the most important tight junction molecules, whose expression is upregulated in the inflamed gut during colitis and is associated with increase in the permeability of the epithelial layer (47). We observed significant downregulation of Claudin-2 expression on treatment with only GILZp or only Py, as well as on combined treatment with both (Fig. 2B). Another important protein involved in protection of the mucus is MUC2, which is the main component of the intestinal mucus layer and usually depleted in IBD (48-51). GILZp, when administered alone and in combination, resulted in the restoration of MUC2 expression, as opposed to Py alone, which did not have this effect (Fig. 2B).

### Restoration of the intestinal burden of clinically important *Candida* and former *Candida* species by GILZp and Py

Recent research has provided evidence for the involvement of the intestinal fungal community, particularly *Candida spp*, including former *Candida* species, in the pathogenesis of IBD (21, 28, 31, 52). To investigate the effects of treatment with Py, GILZp or combination of both (Py+GILZp) on the overall intestinal burden of clinically relevant *Candida* and former *Candida spp*, we conducted RT-qPCR analysis using a previously established protocol (53). First, we validated the assay’s proficiency in identifying our target species, including *C. albicans*, *C. parapsilosis*, *C. dubliniensis*, *C. tropicalis*, *C. sake*, *Debariomyces hansenii* (ex *C. famata*) *Nakaseomyces glabrata (ex C. glabrata*), *Kluyveromyces marxianus (ex C. kefyr*), *Pichia kudriavzevii (ex C. krusei*), *Meyerozyma guillermodii (ex C. guilliermondii*) and *Clavispora lusitaniae* (ex. *C. lusitaniae*) (Fig. 3A) (54). The alignment confirmed the specificity with the target sequences, so we proceeded with the RT-qPCR. Our results show that the administration of DSS significantly elevated the overall intestinal burden of these fungal species compared to the non-colitic group. Importantly, both individual treatments and their combination led to a significant reduction in the same fungal species, and their levels were similar to those observed in non-colitic mice (Fig. 3B).

**FIG. 3.**
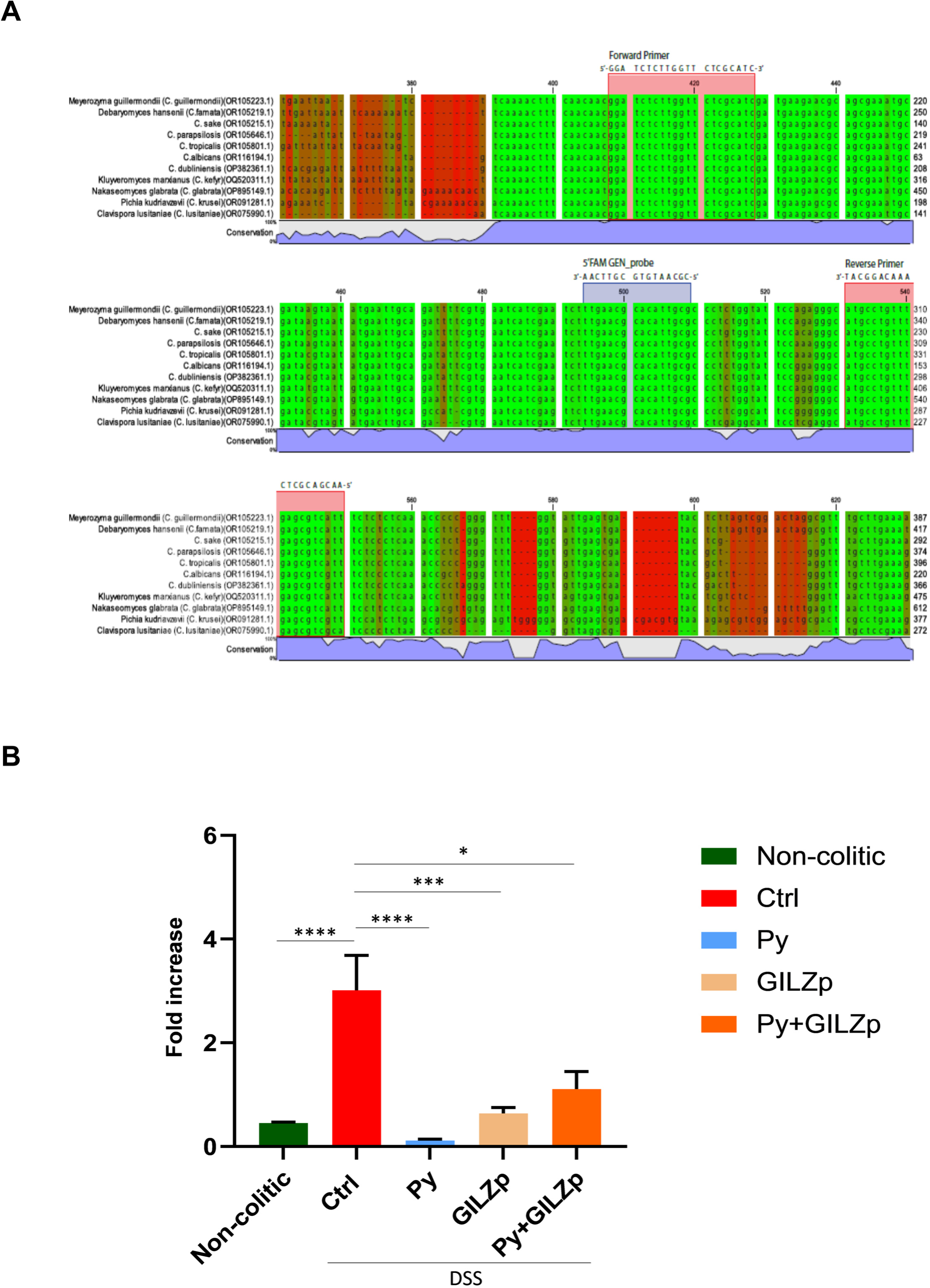
Restoration of *Candida* and former *Candida* species burden to physiological levels in treated colitic mice. **A** Sequence alignment of *Candida* spp. and the main clinically relevant former *Candida spp* was conducted using sequences retrieved from the Nucleotide database. The colors represent the level of homology across species: green indicates perfect homology, and red indicates poor homology. Additionally, the degree of conservation of the sequence between species is indicated below the alignment (represented by the blue bar). The forward and reverse primers (highlighted in red boxes) and the 5′ FAM GEN probe (highlighted in the blue box) were aligned with the sequences. **B** Quantitative RT-PCR for the detection of *Candida* spp. burden in stool samples using a custom TaqMan probe and the ITS1-2 primers to amplify fungal rDNA. Differences between the cycle threshold (Ct) of *Candida* spp. and ITS1-2 were calculated (ΔCts), and data are shown as 2^-ΔCts^. Values are expressed as mean ± SEM in panel B (*n* = 4 per group). * *P* < 0.05, **** *P* < 0.0001.

### Reduction of intestinal fungal dysbiosis in colitic mice by GILZp and Py

Next, we performed mycobiome analysis using high-throughput internal transcribed spacer 1 sequencing of fungal ribosomal DNA in order to characterize the impact of the treatments on the biodiversity of the DSS-associated gut mycobiome and to determine its composition. The total number of raw sequenced reads was 4014891, including 1067582, 821557, and 1651783 reads in the non-colitic, Ctrl, and Py+GILZp/Py/GILZp groups, respectively, with corresponding average values of 213516.4, 136926.17, and 117984.5, respectively. In order to facilitate the comparison of fungal composition between groups and to compensate for biases induced by sequencing library size, bioinformatic analyses were carried out using the rarefaction strategy. Rarefaction curves of the Shannon and Chao1 (Fig. 4A) alpha diversity indexes versus sequencing depth show the goodness of the sequencing data. The curves for each sample (left panel) reached a plateau, thus confirming that the sample sequencing depth was adequate for evaluating both the evenness (the degree of partitioning of taxa abundances in the treatment groups) and the distribution of rare taxa in the samples. In order to evaluate if the available sequencing depth could be used to determine the biodiversity of the treatment groups, alpha indexes were grouped and averaged across treatments (right panels of Fig. 4A). Since the boxplots of both alpha indices of each group are well separated from each other, the acquired dataset with 4000 reads allows for the characterization of both the evenness and the composition of rare taxa in each group of interest. In particular, based on the values of the Shannon indices (Fig. 4A, top right panel), it appeared that the Ctrl and non-colitic groups had the lowest and the highest evenness, respectively. These findings were indicative of the skewed distribution of fungal species in the colitic groups. Furthermore, the curves of the treated groups were close to those of the non-colitic mice, thus indicating that the treatments were able to restore evenness to a point that was close to that of the non-colitic mice. With regard to the Chao1 index (Fig. 4 A, bottom right panel), all the treated groups had lower values than the Ctrl group. This could be due to the fact that, in the period of time corresponding to the completion of the experiment, the number of types of fungi, although higher than that of colitics, has not yet completely re-established itself at values comparable to those of non-colitics in terms of rare fungi. The distribution of alpha indexes (Fig. 4B and Fig. S2) in each group evaluated based on data rarefied with 4000 reads confirms this trend: the alpha diversity indexes revealed that compared to the Ctrl group, on average, the non-colitic (Shannon p.adj = 0.036) and treatment groups had increased evenness, while the non-colitic group had increased richness and the treated groups had reduced richness (Chao1 p.adj = 0.354, Observed_features p.adj = 0.284, and Simpson p.adj = 0.016). All three treated groups showed similar median values for each alpha index, with the average richness being lower than that of the Ctrl group and the evenness being comparable to that of the non-colitic group. Overall, these findings indicate that colitic mice exhibit reduced evenness and richness of their gut fungal community in comparison to non-colitic mice. Both singular and combined treatments showed a clear trend toward restoring the goodness value of alpha diversity, and this indicates their potential to improve the evenness of the fungal community in the gut of colitic mice.

**FIG. 4.**
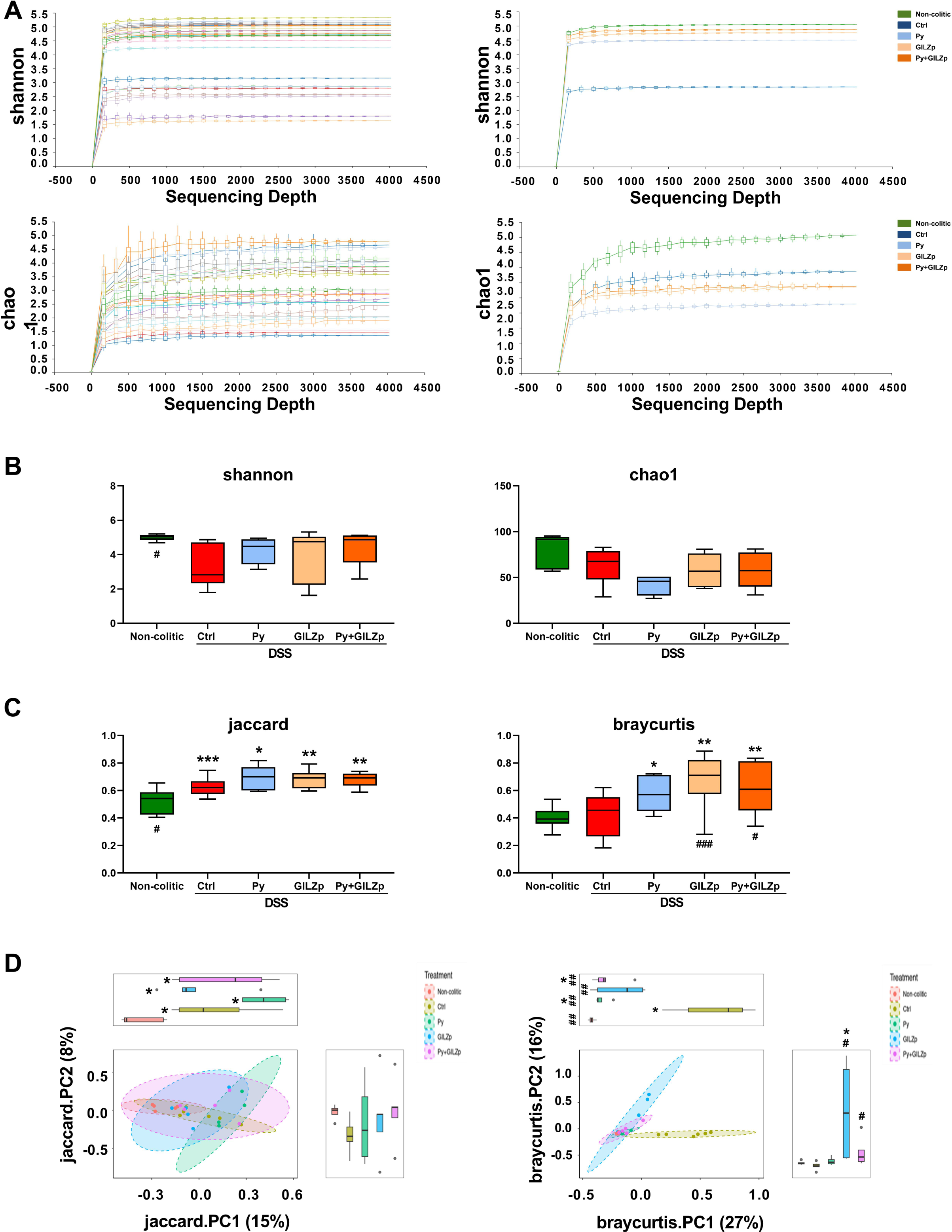
Diversity analyses showing the peculiar fungal composition of non-colitic, control, and treated mice. **A** Rarefaction curves of the alpha diversity indices (Shannon and Chao1) of fecal mycobiota showing the boxplots of each index, evaluated at the sequence variant level, versus sequencing depth. Each boxplot in the left panel represents 50 rarefactions of each sample, which are grouped by treatment in the corresponding right panel, according to the color legend. The curves reach a plateau for all samples, and the treatments have distinct index values when the rarefaction is considered at 4000 reads for each sample. **B** Boxplots of alpha diversity indexes (Shannon and Chao1) of fecal mycobiota evaluated for data rarefied at 4000 reads. Statistical significances were determined using the Wilcoxon multiple-comparison test with FDR adjustment. ^#^*P* < 0.05 for non-colitic *vs* Ctrl group. **C** Boxplots of the beta diversity distances (Jaccard and BrayCurtis) of fecal mycobiota evaluated for data rarefied at 4000 reads. Statistical significances were determined using the Wilcoxon multiple-comparison test with FDR adjustment. The asterisks (*) indicate the significance of differences for the Ctrl or treated group versus the non-colitic group, and the hashtags (^#^) indicate the significance of differences for the non-colitic or treated versus the Ctrl groups. ^#,^\**P* < 0.05, \*\**P* < 0.01, \*\*\**P* < 0.001. **D** PCoA of Jaccard and BrayCurtis distances. The first two PCoA components are represented in a scatter plot and in the marginal boxplots. The percentage of variance explained by each component is shown on each axis. Confidence ellipses assume a multivariate t-distribution. Asterisks (*) indicate the significance of differences for the Ctrl or treated versus non-colitic groups, and hashtags (#) indicate the significance of differences for the non-colitic or treated versus Ctrl groups. ^#,^\**P* < 0.05, ^# #^*P* < 0.01

Permutational multivariate analysis of variance (PERMANOVA) based on non-phylogenetic beta dissimilarity matrices (Jaccard and Bray-Curtis) revealed significant fungal biodiversity in the compositions of groups (Fig 4C). The difference in the Jaccard metric (p.adj=0.024) between the non-colitic and Ctrl mice confirms the difference in their community structure. The qualitative (Jaccard) and quantitative (Bray-Curtis) distances of treated groups present similar values and, thus, further association analyses are needed to evaluate any differences in their fungal composition. Furthermore, since the treated group has significantly higher values than the Ctrl group in terms of both beta diversity indices, the corresponding samples have a more diverse mycobiome both in terms of abundance values and taxa types. According to the Bray-Curtis metric, significant differences were observed between the Py+GILZp and Ctrl groups (p.adj=0.024) as well as between the GILZp and Ctrl groups (p.adj=0.0036). PCoA cluster analysis (Fig. 4D) of beta diversities confirmed the significant compositional diversity of the mycobiome among the different groups. Indeed, the first component of the Jaccard index indicated significant differences in intestinal fungal communities between the non-colitic and Ctrl or treated groups. This indicates that the non-colitic mouse intestine is characterized by specific taxa that are not present in other groups, i.e., a higher degree of richness. Moreover, the first component of the Bray-Curtis metric underlines significant differences in the proportion of fungi between treated mice and non-colitic or Ctrl mice, thus emphasizing the presence of characteristic distributions of taxonomies within each group. The Ctrl group, therefore, presented with significant differences both in the number of types of fungal taxa and in their proportions in comparison with each of the other groups.

Overall, these results indicate that the fungal communities of colitic mice are characterized by richness values that are lower than those in non-colitic mice and that the fungal taxa found in the treated groups are probably also present in the colitic group, but with significant differences in their abundance.

To investigate the composition of the intestinal mycobiota in response to treatment with one compound or combined treatment, taxonomic analysis was performed at various levels, ranging from the phylum to the species level. Our results revealed that the intestinal mycobiota consisted of fungi belonging to the Ascomycota and Basidiomycota phyla across all groups. Specifically, the Ascomycota phylum was found to be the dominant one (Fig. 5A), and the ratio of Ascomycota to Basidomycota was found to be higher in colitic mice than in the non-colitic group mice. Both GILZp and Py+GILZp treatments induced a significant decrease in the Ascomycota/Basidiomycota ratio in comparison to the Ctrl group (P.adj < 0. 05) (Fig. 5A).

**FIG. 5.**
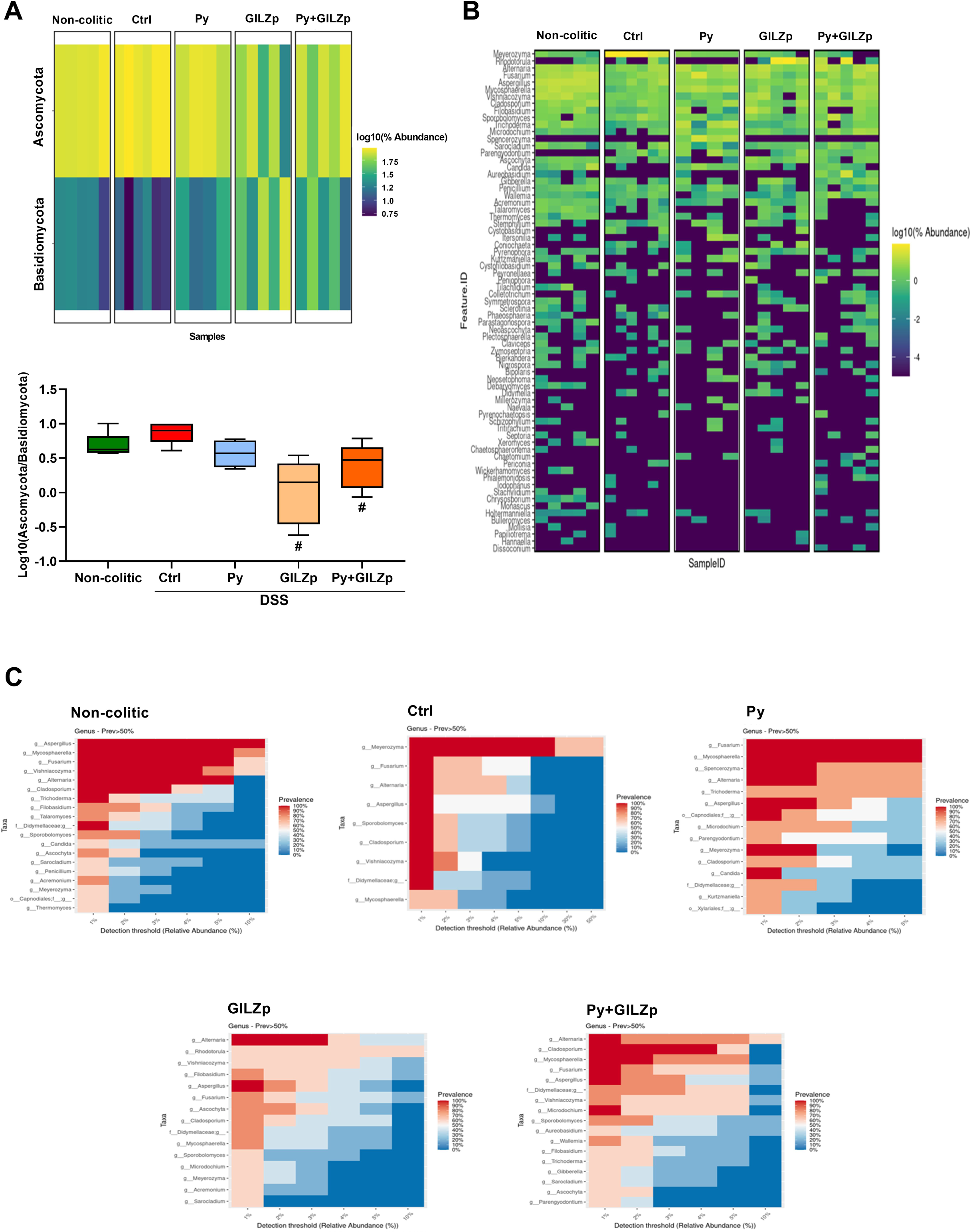
Mycobiome composition (phylum and genus levels) and mycobiome-core (genus levels) analyses**. A** Heatmap depicting the abundances of phyla based on the sequencing data according to groups of interest. Features are sorted from top to bottom by decreasing value of the average abundance calculated over the entire sequencing. Log ratios of abundance between Ascomycota and Basidiomycota phyla for each group. Statistical significances were determined using the Kruskal– Wallis test. ^#^*P* < 0.05 for the GILZp or Py+GILZp group versus the Ctrl group. **B** Heatmap depicting the abundances of genera present based on the sequencing data according to groups of interest. Features are sorted from top to bottom by decreasing value of the average abundance calculated over the entire sequencing. **C** Heatmaps of the mycobiome core of the genera in each sample group, as indicated in each heatmap, versus the per-sample relative abundance threshold (minimum detection threshold = 1%). The core of each group is composed of genera with a relative abundance of at least 1% and with prevalence greater than 50% in the group itself.

At the genus level, heatmap (Fig. 5B) and bar-plot analyses (Fig. S3) showed that the most abundant (on average) genera were *Meyerozyma, Rhodotorula*, *Alternaria*, *Fusarium*, *Aspergillus*, *Mycosphaerella*, *Vyshiacozyma*, *Cladosporium*, *Filobasidium*, and *Sporobolomyces*. Of note, *Meyerozyma* was the dominant genus in the Ctrl group, and its abundance was greatly reduced by singular and combined treatments. According to mycobiome core analysis of each group (Fig. 5C) for fungi with percentage abundance greater than 1%, the following genera showed 100% prevalence in their respective groups, with the minimum value of the percentage abundance of each genera reported in parentheses: *Aspergillus* (10%), *Mycosphaerella* (5%), *Fusarium* (5%), *Alternaria* (5%), *Visnhiacozyma* (4%), *Cladosporium* (3%) and *Trichoderma* (1%) in the non-colitic group; *Meyerozyma* (20%), *Fusarium* (1%), *Alternaria*, (1%), *Aspergillus* (1%), *Sporobolomyces* (1%), *Cladosporium* (1%), and *Visnhiacozyma* (1%) in the Ctrl group; *Fusarium* and *Mycosphaerella* (5%), *Spencerozyma* (2%), *Alternaria* (2%), *Aspergillus* (2%)*, Meyerozyma* (2%), and *Candida* (1%) in the Py group; *Alternaria* (4%) and *Aspergillus* (1%) in the GILZp group; *Cladosporium* (4.5%), *Mycosphaerella* (2.5%), *Fusarium* (1%), *Aspergillus* (1%), and *Microdochium* (1%) in the Py+GILZp group. To identify the fungal taxa driving these community shifts, linear discriminant analysis effect size (LEfSe) analysis was subsequently performed from the phylum to the species level. Fig. 6 shows the results of LEfSe analysis conducted at the genus level through pairwise comparisons of the Ctrl group with the non-colitic (a), Py (b), GILZp (c), or Py+GILZp (d) groups. Among the genera with a prevalence of 100% within the corresponding mycobiome core, highlighted with black arrows, LEfSe analysis demonstrated a consistent and significant association of the *Meyerozyma* genus with the colitic group in all the comparisons conducted (Fig. 6A–D). On the other hand, several genera, such as *Aspergillus*, *Mycosphaerella*, *Fusarium*, *Vishniacozyma*, and *Trichoderma*, showed significant associations with the non-colitic group (Fig. 6B). Our findings indicated that Py and Py+GILZp groups, compared to the Ctrl mice, were enriched with *Mycosphaerella* (Fig 6B and 6D). None of the genera present in the GILZp mycobiome core were significantly associated with the GILZp group (Fig 6C). Further LEfSe analyses of the same pairwise groups at the species level showed that *M. guillermondii* and *M. tassiana* were significantly associated with the Ctrl group and the non-colitic or Py+GILZp group, respectively. These results suggest that the *Meyerozyma* genus could be one of the key microbial genera characterizing the gut fungal community in DSS-induced colitis. Furthermore, the reduction in its abundance may contribute to the shift of the intestinal fungal community toward eubiosis, thereby contributing to the observed anti-colitic effects with both singular and combination treatments.

**FIG. 6.**
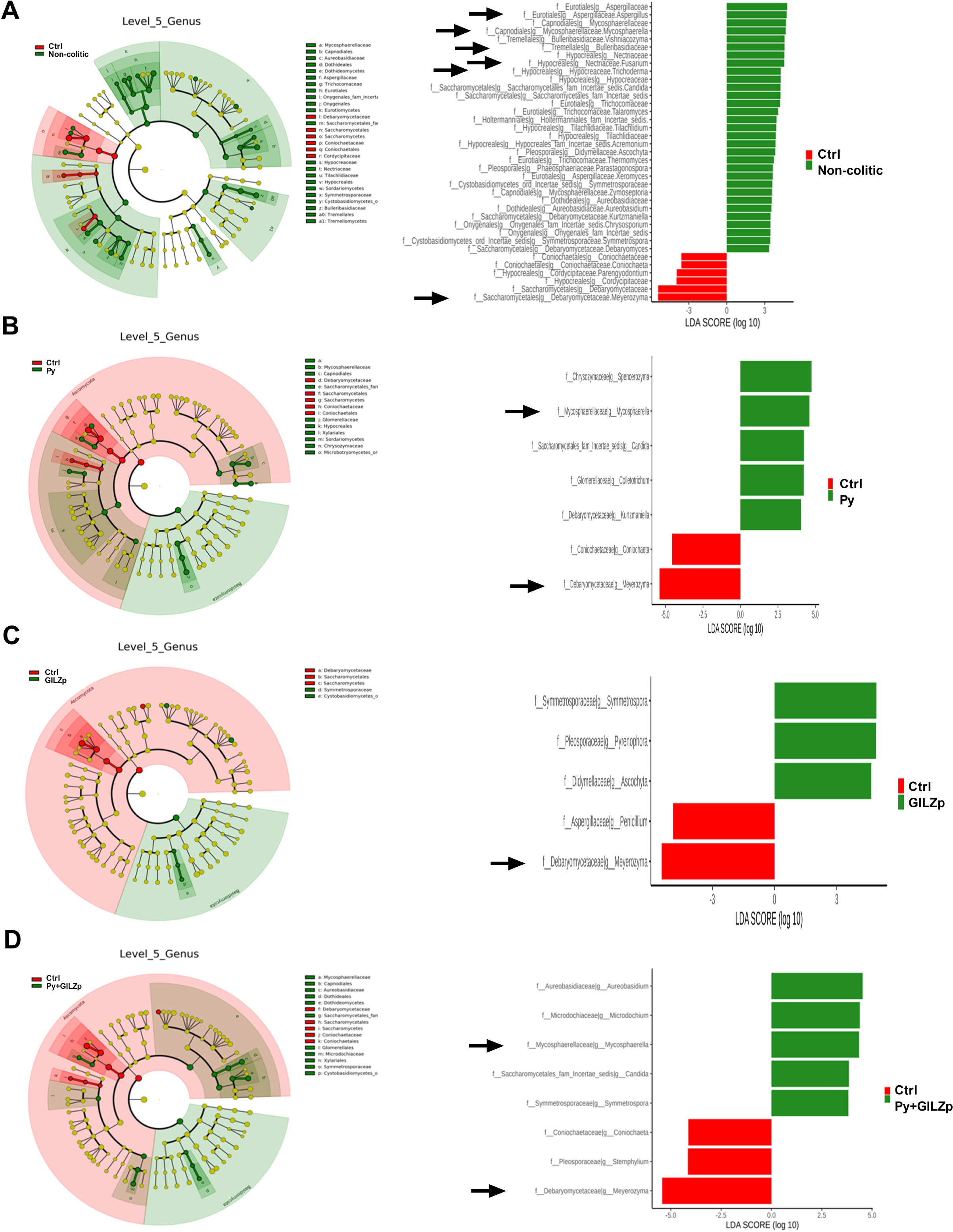
Differential association analysis of genera in each treated or non-colitic group in comparison with the Ctrl group. **A–D** Cladograms and histograms showing taxonomies, up to the genus level, that were significantly (*P* < 0.05) associated with one of the two groups considered, as indicated in the corresponding color legend. The yellow circles in the cladogram indicate non-significant taxa. The black arrows in the histograms emphasize the genera belonging to the mycobiome core of the group itself, from among the genera significantly associated with each group.

## DISCUSSION

Despite the availability of several classes of drugs for the relief of symptoms and reduction of inflammation in IBD, many patients either do not respond to treatment or experience a loss of response over time and, often, need to switch to another class of drugs. However, untill date, no definitive cure has yet been found for IBD. Additionally, long-term treatments with some efficacious drugs, such as glucocorticoids and other immunosuppressants, lead to serious adverse effects, including metabolic alterations and increased risk of infections and malignancies (55, 56). GILZ is a protein that mediates several anti-inflammatory effects of glucocorticoids, and previous studies on GILZp have reported promising results for the treatment of IBD (6, 8, 57-60). Considering that GILZ is also involved in the pathophysiology of IBD, as demonstrated in both preclinical models of colitis and in patients, in this study, we investigated the effects of combining GILZp with a potential prebiotic yeast extract (Py) in a mouse model of colitis. In our initial experimental model of colitis, the mortality observed in the Ctrl mice was considerably prevented by Py in a dose-dependent manner. Based on this finding, one hypothesis that explains the observed effect is that Py could play a protective role in the mucosa, to mechanically prevent the disruption of the mucosal barrier. However, the treated mice in this study showed symptoms of IBD, such as loss of body weight and a DAI score that was similar to that of Ctrl mice, at the end of the experiment. Alternatively, similarly to prebiotics, Py may favor the growth of beneficial microbiota, and this may ultimately relieve the symptoms of colitis. Interestingly, treatment with only GILZp led to the survival of colitic mice, with the molecular events underlying this beneficial effect only partially overlapping with those of Py. In particular, GILZp, by acting selectively on MUC2, improved mucus production, thus helping in the restoration of the mucus layer. In contrast, Py acted mainly on pro-inflammatory cytokines, by either inhibiting their production or reducing their expression. Of note, both GILZp and Py reduced Claudin-2 expression: this implies that they both play a role in improving barrier integrity and alleviating the symptoms of the disease (61). Overall, both GILZp and Py treatments promoted the survival of mice and led to milder disease by acting on different and overlapping molecular targets. One limit of this study was the short follow-up time, since a longer experimental duration can lead to potential differences between treated and Ctrl groups in the final DAI score.

Recent studies have suggested that intestinal fungal dysbiosis potentially contribute to the development and persistence of IBD. Therefore, it is important to investigate the fungal microbiome to gain a comprehensive understanding of the complex interactions between microbial communities and the immune system in this context (21, 22, 31, 33, 52, 61-63). We therefore aimed to examine the impact of administering Py, GILZp, and a combination of both on the composition of the intestinal fungal community. Initially, we conducted RT-qPCR analysis of fecal samples obtained from non-colitic, colitic, and treated colitic mice, to determine the intestinal load of clinically important *Candida* and former *Candida* species, namely, *Debaryomyces hansenii, Meyerozyma guillermondii,* and *Clavispora lusitaniae*, previously known as *Candida famata, Candida guillermondii*, and *Candida lusitaniae*, respectively (54). In accordance with previous findings (33, 64, 65), our results revealed a significant increase in their overall intestinal load in colitic mice compared to the non-colitic mice. The administration of Py, GILZp, or their combination restored their intestinal load and brought it up to levels similar to those observed in non-colitic mice. We cannot exclude the direct effect of Py or GILZp on *Candida* spp., but it is plausible that this outcome is the result of several indirect effects that are primarily driven by stimulation of the growth of beneficial intestinal bacteria. Indeed, as a prebiotic (36), Py is able to induce the selective proliferation of bacteria, such as *Lactobacillus* and *Bifidobacterium* species (https://procelys.com/living-products/nucel/). Both species are well known for their capacity to reduce the overgrowth of opportunistic pathogens, such as *Candida* spp., by multiple mechanisms including competition for adhesion sites and nutrients, secretion of antimicrobial compounds, reduction of inflammatory cytokines such as IL-1α, IL-1β, IL-6, IL-12, TNF-α, and IFN-γ), improvement of intestinal barrier integrity, and production of butyrate and short-chain fatty acids (66-68). We hypothesize that, also, the effect of GILZp is mediated by its action on beneficial bacteria, but unlike the effect of Py, this effect of GILZp is exerted in an indirect manner. Indeed, we recently demonstrated that GILZp can restore an optimal environment for colonization of health-promoting bacteria by acting on the mechanisms of control of the integrity and functions of the epithelial barrier (9).

Our analysis of the fungal community revealed that colitic mice were characterized by reduced alpha-diversity and by a distinct taxonomic composition in comparison to non-colitic mice. This is in line with previous findings (21, 28, 52). Py and GILZp, when administered alone and in combination, were effective in reducing DSS-induced fungal dysbiosis and promoting a fungal composition that was similar to that of non-colitic mice. Further, consistent with previous studies (52, 69-72), our results indicate that the composition of the intestinal mycobiome in both non-colitic and colitic mice is dominated by fungi from the Ascomycota and Basidiomycota phyla. In particular, Ascomycota consistently appeared to be the most abundant phylum in all the groups. Dysbiosis of the Ascomycota and Basidiomycota phyla in IBD, such as ulcerative colitis, is complex and can vary among individuals. Currently, the data regarding the Ascomycota*/*Basidiomycota ratio in patients with ulcerative colitis compared to non-colitic subjects are discordant. Some studies have reported that their ratio is not significantly different compared to non-colitic subjects (30, 52). In contrast, other studies have reported an increase in their ratio (73, 74) and one study reported a decrease in the ratio in patients with ulcerative colitis (21). Our results show a marked increase in the Ascomycota to Basidiomycota abundance ratio in 80% of colitic mice compared to the non-colitic controls, thus supporting previous data indicating a strong association between the abundance of Ascomycota and ulcerative colitis. Combined treatment with GILZp and Py significantly reversed the imbalance between Ascomycota and Basidiomycota. Based on the increased abundance of Ascomycota observed in colitic mice, and in line with RT-qPCR results, an in-depth analysis of the mycobiota taxonomic composition showed that the DSS group was dominated by the *Meyerozyma* genus (phylum Ascomycota, family *Debariomycetaceae*, species *M. guillermondii* species, formerly *Candida guillermondii*). Thus, it could be one of the main microbial taxa involved in the pathogenesis of ulcerative colitis. Both GILZp and Py, when administered individually or in combination, resulted in a significant reduction in *Meyerozyma* abundance compared to the Ctrl group. Although the mechanistic basis for this effect requires further study, our data suggest that a reduction of gut colonization by *Meyerozyma* spp. could contribute to a positive outcome in patients with ulcerative colitis. Furthermore, our results showed that the *Mycosphaerella* genus was one of the predominant fungal taxonomic units in the intestinal fungal community of non-colitic mice (specifically *M. tassiana*, previously known as *Davidiella tassiana*, the sexual form of *Cladosporium herbarum*, which belongs to phylum Ascomycota, family *Mycosphaerellaceae*), whereas it was not significantly associated with colitic mice. Interestingly, this fungus showed a significant association with both the Py and Py+GILZp groups. Thus, there might be a potential link between *Mycosphaerella* and anti-colitic effects mediated by Py. Understanding the role of specific fungal species, such as *M. guillermondii* and *M. tassiana*, in restoring mycobiome balance is crucial for developing potential therapeutic strategies to promote gut health and alleviate gastrointestinal disorders.

### Conclusions

Collectively, our data provide compelling evidence that GILZp and Py cotreatment, by enhancing intestinal barrier function and reducing inflammation and fungal dysbiosis, exerts a protective effect in a mouse model of ulcerative colitis. These findings represent a promising starting point for further investigation into the potential use of their combination as a novel therapeutic strategy for managing IBD.

## Data availability

The metagenomics datasets generated during the current study have been deposited in the Sequence Read Archive (SRA) (https://www.ncbi.nlm.nih.gov/sra) under BioProject ID number PRJNA1021184. All the others generated during the current study are available upon reasonable request.

## Supplemental Material

Extended version of material and methods. Supplementary figures, Fig.S1, Fig.S2, and Fig.S3.

